# Cross-task perceptual learning of object recognition in simulated retinal implant perception

**DOI:** 10.1101/360669

**Authors:** Lihui Wang, Fariba Sharifian, Jonathan Napp, Carola Nath, Stefan Pollmann

## Abstract

The perception gained by retina implants (RI) is limited, which asks for a learning regime to improve patients’ visual perception. Here we simulated RI vision and investigated if object recognition in RI patients can be improved and maintained through training. Importantly, we asked if the trained object recognition can be generalized to a new task context, and to new viewpoints of the trained objects. For this purpose, we adopted two training tasks, a naming task where participants had to choose the correct label out of other distracting labels for the presented object, and a discrimination task where participants had to choose the correct object out of other distracting objects to match the presented label. Our results showed that, despite of the task order, recognition performance was improved in both tasks and lasted at least for a week. The improved object recognition, however, can be transferred only from the naming task to the discrimination task but not vice versa. Additionally, the trained object recognition can be transferred to new viewpoints of the trained objects only in the naming task but not in the discrimination task. Training with the naming task is therefore recommended for RI patients to achieve persistent and flexible visual perception.

## Introduction

A growing number of people worldwide are suffering from photoreceptor degeneration in the retina, which characterizes diseases such as retinitis pigmentosa (RP) and age-related macular degeneration (AMD), and causes gradual vision loss (Bourne et al., 2017; Congdon et al., 2004). In an effort to restore vision to patients who suffer from photoreceptor diseases, great progress has been made in the development of visual prostheses (Fine and Boynton, 2015). One of these technologies is the retinal implant (RI), a photoelectric device which stimulates the remaining neurons in the retina to evoke action potentials that can be transmitted to the visual cortex to form visual percepts (Shepherd et al., 2013; Zrenner, 2002).

Clinical trials with RI systems that have obtained regulatory approval for clinical treatment (Argus II system, Ho et al., 2015 and Alpha IMS system, Zrenner et a., 2011) report that the patients were able to discriminate orientation and motion direction (Ho et al., 2015), and even read large letters (Zrenner et al., 2011). Despite these promising results, the perception gained by RI is still limited (Shepherd et al., 2013), and there is a large variation of the gained perception according to the few clinical reports. Such limitations raise the question whether, and to what extent, RI perception can be improved through training.

It is well documented that training on a visual task can significantly improve visual perception. These improvements, due to visual perceptual plasticity, can persist for months or even years (Goldstone, 1998). Training on a visual task can significantly strengthen visual perception in both healthy subjects and patients with visual deficits (Ooi et al, 2013; Polat et al., 2004). Visual perceptual plasticity can be gained from the familiarity of a particular task (task-dependent), or from boosted representation of the trained feature irrelevant of the task (task-independent, Watanabe and Sasaki, 2015). The task-dependent plasticity constrains the improved visual performance to a particular task. Therefore, it is important to achieve task-independent perceptual plasticity for RI users such that the improved visual perception can generalize to other contexts.

The goal of our study was to develop a training paradigm that can help RI patients to make optimal use of their newfound vision. To reduce unnecessary testing burden for patients, we conducted simulation experiments mimicking the limited vision of the patients with a subretinal implant (Perez Fornos et al., 2008). Such simulated RI-vision was realized by reducing spatial resolution, field of view and implementing specific distortions in pictures shown to observers with intact vision (Beyeler et al., 2017; Chen et al., 2009). Specifically, we focused on the visual ability of recognizing everyday objects. With the large amounts of variants for each object and the different retinal representations of the same individual object given by different viewpoints, we were able to test the flexibility of the trained visual ability. Here we investigated if a perceptual learning regime can improve object recognition in simulated RI-vision, and if the improved object recognition can be transferred to a different task. For this purpose, we included two different tasks, a naming task where participants have to choose the correct label out of other distracting labels for the presented object, and a discrimination task where participants have to choose the correct object out of other distracting objects to match the presented label (Figure 1B). Both tasks required the discrimination of the correct object representation from a set of competing object representations. They differed in that either one visually presented object needed to be compared to memory traces of visual objects (in the naming task) or one visual object memory trace needed to be compared to a set of object pictures (in the discrimination task). In principle, we expected to observe improvement in object recognition irrespective of the particular training type. However, the discrimination task lends itself to a feature-based comparison of object pictures, whereas the naming task emphasizes a more holistic comparison of the stimulus with object memory traces (Song et al., 2010). Given the reduction of distinctive features in RI vision, a holistic strategy might be more successful than a feature-based strategy.

**Figure 1.**
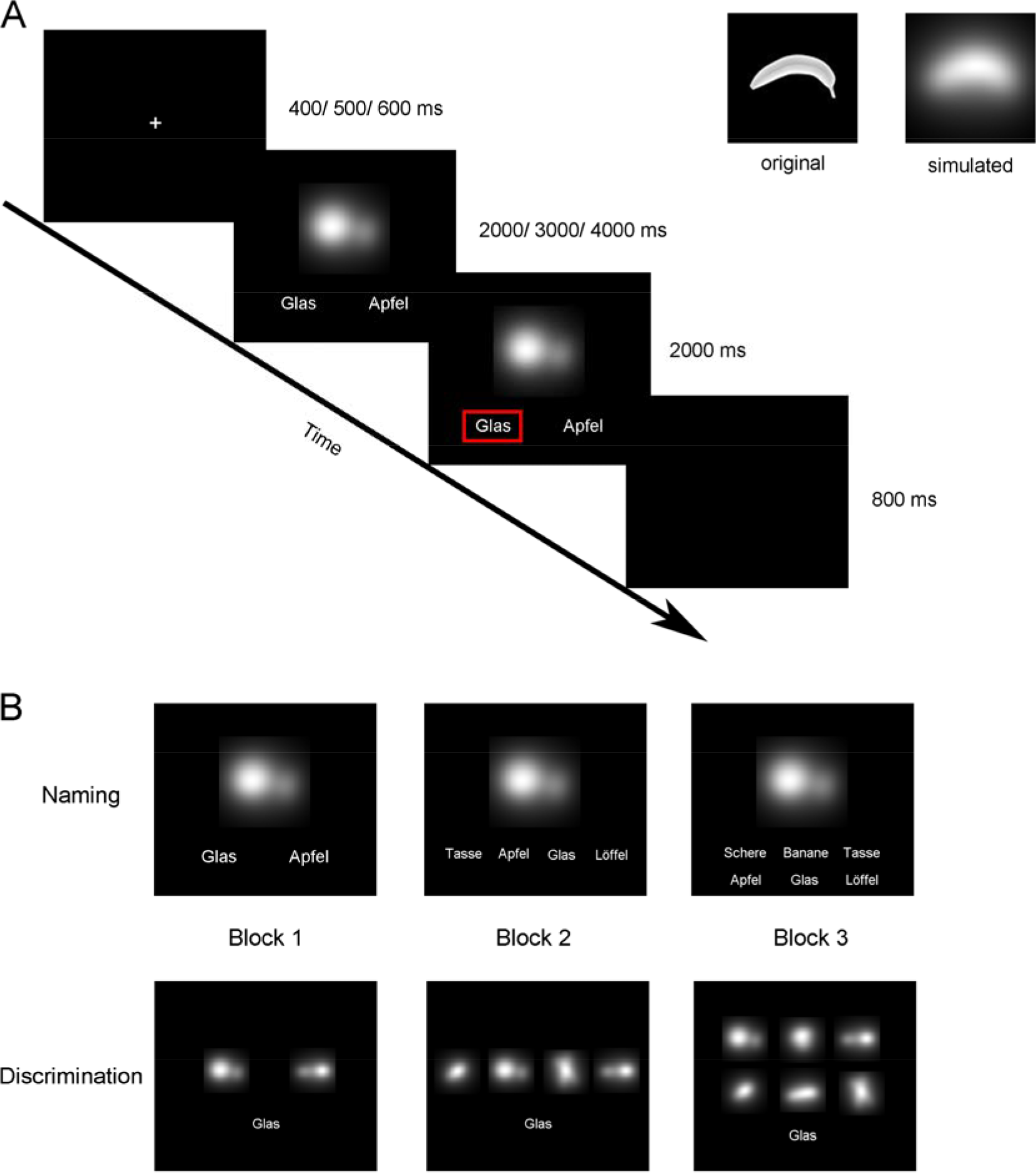
**A:** an example trial in the naming task (left), the original and the simulated picture of a banana (right). **B**: The naming task (upper) and the discrimination task (lower) in Experiment 1 where the number of the alternative choices increased over the three blocks. In Experiment 2, the number of alternative choices was always 4.

We hypothesized that if object recognition training can transfer across tasks, the improved performance in one task would persist, or could be even further boosted, in the other task. Taking advantage of the different viewpoints of everyday objects, we also asked whether the object recognition of trained viewpoints can transfer to new, untrained viewpoints. Moreover, it has been shown that learning fine visual discriminations can be speeded up by insertion of easy trials into the training (Ahissar and Hochstein, 1997; Maertens and Pollmann, 2005; Rubin et al., 1997). Once a feature difference has been detected, it will lower the discrimination threshold for this feature in the future. Based on these findings, we manipulated the task difficulty in Experiment 1 by varying the number of alternative choices in the hope that the easier trials at the beginning of the task could boost the learning process. To test if such manipulation made a difference in producing the training effect, the task difficulty in terms of the alternative choices were controlled in Experiment 2. Additionally, the persistence of the improved object recognition was further tested in Experiment 2 by including a post-test which took place one week after the training session.

## Experiment 1

### Methods

#### Participants

Thirty-three healthy students participated in Experiment 1, with 16 of them (8 females, Age: 18 - 31 years old) randomly assigned to Experiment 1A, and the other 17 (12 females, Age: 18 - 26 years old) assigned to Experiment 1B. All participants had normal or corrected-to-normal vision, and all of them were German native speakers. This experiment was conducted in accordance with the Declaration of Helsinki and was approved by the local ethics review board. Written consent form was obtained from each of the participants prior to the experiment. None of them had been exposed to the simulated pictures before the experiment.

#### Stimuli and Design

In both experiments, stimuli were simulated object pictures using Pulse2percept software (Beyeler et al., 2017), with the technical parameters of the subretinal Alpha IMS system (Retina Implant AG, Tübingen; Zrenner et al., 2011), i.e. each picture had a resolution of 1369 stimulating electrodes in an area covering a visual field of 8° visual angle. The likelihood of axonal stimulation was set  = 0.1, because of the subretinal stimulation of the bipolar cell layer. Eight different objects (Apple, Banana, Bottle, Cup, Glass, Spoon, Scissors, and Toothpaste) from different viewpoints were simulated, rendering 64 pictures in total.

There were two tasks in each of the two experiments: a naming task (Figure 1B, upper) and a discrimination task (Figure 1B, lower). In the naming task, one object picture was presented at the center of the screen, below which 2, 4, or 6 German words referring to object names were presented. Participants were asked to choose the correct name for the object picture by clicking the left button of the mouse. In the discrimination task, one object name was presented at the lower visual field of the screen, above which 2, 4, or 6 object pictures were presented. Participants were asked to choose the correct picture matching the name by clicking the left button of the mouse. Each task had an increasing difficulty over three consecutive blocks in the way that there were two alternative choices in Block 1, 4 in Block 2, and 6 in Block 3. In Experiment 1A, the naming task was followed by the discrimination task; in Experiment 1B, the order of the two tasks was reversed.

#### Procedures

Each trial began with a white fixation cross at the center of a black screen for a duration randomly selected from 400/ 500/ 600 ms (Figure 1A, left). Then the task frame, which contained the picture(s) and name(s), was presented until the first mouse click or the time limit was reached. The time limit was 2s when there were 2 alternative choices (Block 1), 3s when there were 4 alternative choices (Block 2) and 4s when there were 6 alternative choices (Block 3). Mouse click beyond this time limit was identified as a response omission for the current trial. After the task frame, a feedback frame was presented for 2s. The feedback frame was the same as the task frame except that the correct picture or name was marked by a red box. The inter-trial-interval was a blank screen of 800ms. Participants were asked to respond as fast and accurately as possible.

For both experiments, there were 64 trials in each block. Each picture appeared as the target picture only once in each block. In each block, the 64 trials were presented in a pseudorandom order in the way that 8 objects from 4 different viewpoints were followed by the same 8 objects from 4 additional different viewpoints, rendering 2 mini-blocks in each block and 12 mini-blocks in total. This arrangement was chosen to investigate if the training on the first 4 viewpoints could be transferred to the untrained new viewpoints, particularly in Block 1 where all of the pictures were presented for the first time. Thus, each picture appeared as the target picture three times in total in each task. Within each mini-block, the distracting picture(s) or name(s) in each trial was randomly selected from other object pictures or names. Each picture/ name appeared as one of the distractors with equal probability. There was a 1min break after each block, and a 3min break between the two tasks. Prior to each task, 6 example trials with 2 alternative choices were provided. In these example trials, the original picture (without simulation) of different objects (Clock, Shoe, Cherry, Headphone, Brush, and Blob) was presented, to prevent participants’ exposure to the simulated pictures.

#### Statistical analysis of data

For each participant, the raw accuracy in each session was calculated as the percentage of trials with a correct response. The theoretical chance-level accuracy (50% in Block 1, 25% in Block 2 and 16.7% in Block 3) was then subtracted from the raw accuracy, rendering the above-chance accuracy (ACA) in each block, such that the accuracies from different blocks could be compared. A 2 (task: naming vs. discrimination) * 3 (block order: 1, 2, vs. 3) repeated-measures ANOVA was conducted on the ACA.

To show the learning curve over the repetition of the pictures across two tasks, the ACA was calculated for each mini-block and shown as a function of the 12 mini-blocks.

The ACAs for the last mini-block in the first task (6^th^ mini-block in total) and the ACAs for the first mini-block in the second task (7^th^ mini-block in total) were compared with a paired-*t* test. If the ACA was significantly lower in the 7^th^ mini-block, the transfer hypothesis was rejected. Alternatively, transfer from the first task to the second task was inferred from one of two outcomes: 1) the ACA in the 7^th^ mini-block was higher than the ACA in the 6^th^ mini-block; 2) the ACA in the 7^th^ mini-block was equivalent to the ACA in the 6^th^ mini-block. Therefore, in case of a non-significant t-test, we calculated a directional Bayes factor BF0- (Rouder et al., 2012; Wagenmakers et al., 2018) using JASP (Wagenmakers et al., 2018b) to quantify the extent to which the null hypothesis was supported. The B_0-_ is the variant of Bayesian null hypothesis testing (often denoted B_01_) where the ‘-‘sign denotes the direction of the alternative hypothesis, given a one-sided test; see supplementary Figure 1 for an example of this type of BF analysis. Here, the BF_0-_ describes the relative probability of the data under the null hypothesis that the ACA in mini-block 7 was equal or higher than the ACA in mini-block 6, relative to the alternative hypothesis that the ACA in mini-block 7 was lower than in mini-block 6. Per convention, a BF > 3 is taken as moderate evidence for the tested hypothesis (Wagenmakers et al., 2018b).

Moreover, the ACAs in the first mini-block in the first task (1^st^ mini-block in total) and the first mini-block in the second task (7^th^ mini-block in total) were also compared with a paired t-test. A significant ACA increase in this comparison was taken as an additional indicator of transfer from the first to the second task, in addition to an ACA increase or a null effect between the 6^th^ mini-block and the 7^th^ mini-block. By contrast, no increase from the 1^st^ mini-block to the 7^th^ mini-block would speak against transfer. The ACAs for the two mini-blocks within Block 1 were further compared using a paired-*t* test, to show if the trained object recognition could be transferred to untrained viewpoints.

### Results

#### Experiment 1A: Naming followed by Discrimination

The 2*3 ANOVA showed a main effect of task, *F*(1,15) = 23.68, *p* < 0.001, *η*^2^ = 0.612, indicating higher ACA in the discrimination task (29.9%) than in the naming task (23.4%), and a main effect of block, *F*(2, 30) = 20.83, *p* < 0.001, *η*^2^ = 0.581. This main effect was due to increasing accuracy over the three blocks (mean ACAs: 20.0%, 29.1%, 30.8%), as revealed by a linear trend, *F*(1, 15) = 23.39, *p* < 0.001, *η*^2^ = 0.609 (Figure 2A, upper). There was an interaction trend between task and block order, *F*(2, 30) = 2.52, *p* = 0.097, *η*^2^ = 0.144. A paired *t* test yielded no evidence that the ACA for the last mini-block of the naming task (6^th^ mini-block in total, 27.9%) was higher than the ACA for the first mini-block of the discrimination task (7^th^ mini-block in total, 24.0%), *t*(15) = 1.20, *p* = 0.124 (one-sided). The BF analysis showed that B_0-_ = 1.231, indicating that the null hypothesis (‘the ACA in the 7^th^ mini-block was equal to or higher than the ACA in the 6^th^ mini-block’) was 1.231 times more likely to be true than the alternative hypothesis (‘the ACA in the 7^th^ mini-block was lower than the ACA in the 6^th^ mini-block’). The B_0-_ here may not be taken as strong evidence (e.g., > 3) to accept either the null hypothesis or the alternative hypothesis. However, the ACA in the 7^th^ mini-block was higher than the ACA in the 1^st^ mini-block (10.9%), *t*(15) = 3.92, *p* = 0.001 (Figure 3A, left). These results indicated that the improved object recognition in the naming task to some extent, if not fully, transferred to the discrimination task. The reliability of the task transfer observed here would be further tested in Experiment 2. Moreover, the paired-t test on the ACAs within Block 1 showed that the ACAs for the second mini-block (18.8%) was higher than the ACAs for the first mini-block (10.9%), *t*(15) = 2.32, *p* = 0.035, suggesting that the object recognition of the trained viewpoints transferred to untrained, new viewpoints.

**Figure 2.**
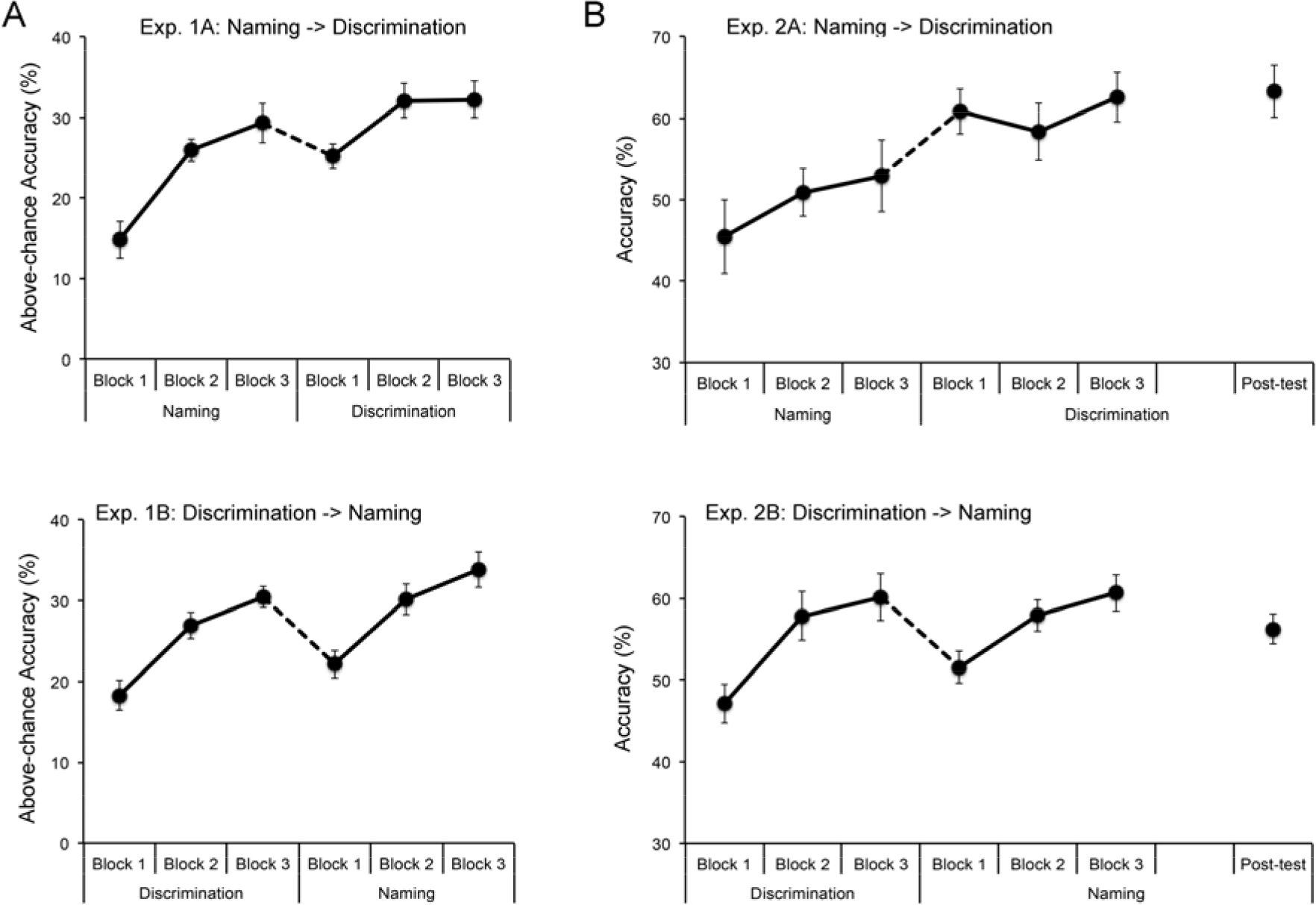
**A:** above-chance accuracies with standard errors are shown as a function of block order and task type in Experiment 1A (upper) and Experiment 1B (lower). **B**: accuracies with standard errors are shown as a function of block order and task type in Experiment 2A (upper) and Experiment 2B (lower). For the post-test which was completed one week after the training, the task was the same as the second task during training. The dashed line indicates a change in task type.

**Figure 3.**
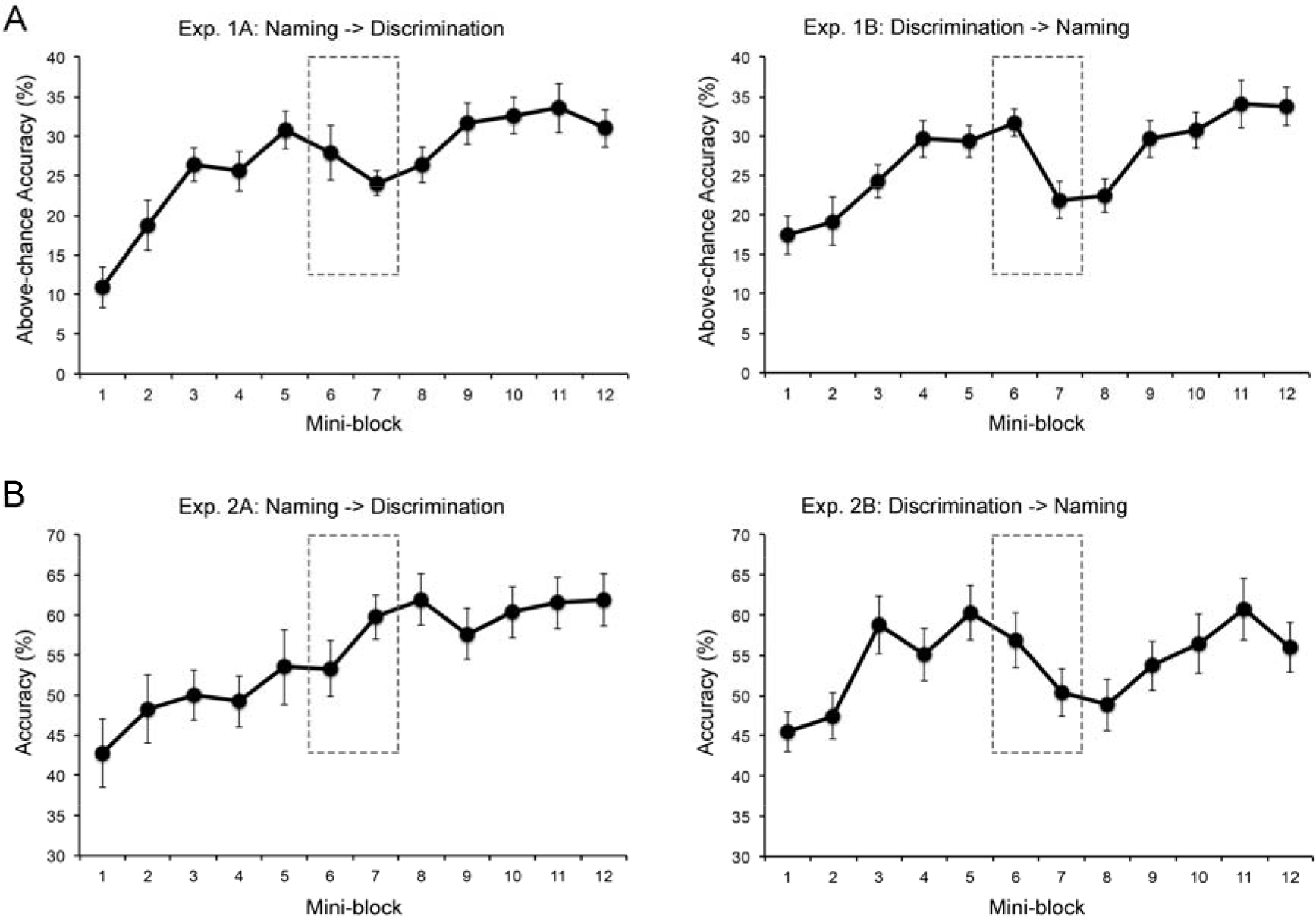
**A:** above-chance accuracies with standard errors are shown as a function of the 12 mini-blocks in Experiment 1A (left) and Experiment 1B (right). **B**: accuracies with standard errors are shown as a function of the 12 mini-blocks in Experiment 2A (left) and Experiment 2B (right). The dashed square indicates the above-chance accuracy/ accuracy in the last mini-block of the first task, and that in the first mini-block of the second task, from which a transfer between tasks can be inferred.

#### Experiment 1B: Discrimination followed by Naming

The 2*3 ANOVA showed a main effect of task, *F*(1,16) = 4.85, *p* = 0.043, *η*^2^ = 0.233, indicating higher ACA in the naming task (28.7%) than in the discrimination task (25.2%), and a main effect of block order, *F*(2, 32) = 36.04, *p* < 0.001, *η*^2^ = 0.693. This main effect was due to increasing accuracy over the three blocks (20.2%, 28.5%, 32.2%), as shown by a linear trend, *F*(1, 16) = 47.39, *p* < 0.001, *η*^2^ = 0.748 (Figure 2A, lower). However, the interaction between task and block order did not reach significance, *F* < 1.

There was a drop of ACA from the 6^th^ mini-block (31.7%) to the 7^th^ mini-block (21.9%), *t*(16) = 3.55, *p* = 0.001 (one-sided), while the ACA in the 1^st^ mini-block (17.5%) did not differ from the ACA in the 7^th^ mini-block (21.9%), *t*(16) = 1.22, *p* = 0.241. These results indicated the absence of the transfer from the discrimination task to the naming task (Figure 3A, right). The transfer from trained viewpoints to untrained viewpoints was also absent, as revealed by the null effect between the first two mini-blocks of the discrimination task (17.5% vs. 19.1%), *t* <1.

## Experiment 2

### Methods

#### Participants

Another 32 healthy students who have never been exposed to the simulated pictures participated in Experiment 2, with half of them randomly assigned to Experiment 2A (10 females, 19 – 29 years old) and the other half assigned to Experiment 2B (10 females, 19 – 28 years old). All participants had normal or corrected-to-normal vision, and all of them were German native speakers. This experiment was conducted in accordance with the Declaration of Helsinki and was approved by the local ethics review board. A written consent form was obtained from each of the participants prior to the experiment. Two participants from each group did not show up in the post-test, and the analysis on the persistency of the training effect was hereby conducted on the remaining 14 participants (Experiment 2A: 9 females, 19 – 29 years old; Experiment 2B: 8 females, 19 – 28 years old).

#### Stimuli and Design

Stimuli and design in Experiment 2 were the same as in Experiment 1 except that there were always 4 alternative choices across all blocks, and the time limit for accepting response was 3s. There was an additional block of post-test, which took place one week after the training. The task in the post-test was the same as the second task during training, to avoid contamination from the other task.

#### Statistical analysis of data

Statistical analysis was equivalent to Experiment 1 except that the raw accuracy (ACC) rather than ACA was compared because of comparable task difficulties over the blocks. To investigate whether and to which extent the training effect persists after one week, a repeated measures ANOVA with post-hoc comparisons was conducted on the accuracies in the first training block, the last training block and the post-test. Similar to the one-sided *t*-tests on task transfer, the null hypothesis for the persistence was ‘the accuracy in the post-test was equal to or higher than the accuracy in the last training block’ while the alternative hypothesis was ‘the accuracy in the post-test was lower than the accuracy in the last training block’. In case a non-significant *p* value was observed, B_0-_ was accordingly calculated to determine the extent to which the null hypothesis is reliable when a non-significant *p* value was observed.

### Results

#### Experiment 2A: Naming followed by Discrimination

The 2 * 3 ANOVA on raw accuracy showed a main effect of task, *F*(1, 15) = 27.55, *p* < 0.001, *η*^2^ = 0.647, indicating increased accuracy in the discrimination task (60.5%) relative to the naming task (49.5%) (Figure 2B, upper). However, the main effect of block, *F*(2, 30) = 2.34, *p* = 0.113, and the interaction, *F*(2, 30) = 2.45, *p* = 0.104, did not reach significance. Although the main effect of block and the interaction were too weak to reach statistical significance, a separate ANOVA showed a main effect of block only for the naming task, *F*(2, 30) = 3.47, *p* = 0.044, *η*^2^ = 0.188, which was due to a linear increasing trend over blocks (45.5%, 49.6%, 53.4%), *F*(1, 15) = 6.22, *p* = 0.025, *η*^2^ = 0.293, but not for the discrimination task, *F* <1. The accuracy did not drop from the 6^th^ mini-block (53.3%) to the 7^th^ mini-block (59.8%), *t*(15) = 1.73, *p* = 0.948 (one-sided). The BF analysis showed that B_0-_ = 9.218, indicating that the null hypothesis (‘the accuracy in the 7^th^ mini-block was equal to or higher than the accuracy in the 6^th^ mini-block’) was 9.218 times more likely to be true than the alternative hypothesis (‘the accuracy in the 7^th^ mini-block was lower than the accuracy in the 6^th^ mini-block’). Moreover, the accuracy in the 7^th^ mini-block was higher than the accuracy in the first mini-block (42.8%), *t*(15) = 3.65, *p* = 0.002. These results indicated a transfer between the two tasks (Figure 3B, left). There was also a transfer from trained viewpoints to untrained viewpoints, as shown by the increased accuracy from the first mini-block (42.8%) to the second mini-block of the naming task (48.2%), *t*(15) = 2.41, *p* = 0.029.

For the persistency of the training effect, the ANOVA showed a significant main effect, *F*(2, 26) = 18.31, *p* < 0.001, *η*^2^ = 0.585, with the accuracies in the last training block (62.6%) and the post-test (63.3%) both higher than the accuracy in the first training block (45.4%), *p* = 0.003 and *p* < 0.001, respectively (with Bonferroni corrections), whereas the accuracy in the last training block did not differ from the accuracy in the post-test, *p* > 0.9. The BF analysis comparing the accuracy in the training block and the accuracy in the post-test showed that B_0-_ = 4.739, indicating that the null hypothesis (‘the accuracy in the post-test was equal to or higher than the accuracy in the last training block’) was 4.739 times more likely to be true than the alternative hypothesis (‘the accuracy in the post-test was lower than the accuracy in the last training block’) (Figure 2B, upper). These results suggested that the improved object recognition by training can last at least for a week.

#### Experiment 2B: Discrimination followed by Naming

The overall accuracy in the discrimination task did not significantly differ from the overall accuracy in the naming task, *F* < 1. However, there was a significant main effect of block, *F*(2, 30) = 28.33, *p* < 0.001, *η*^2^ = 0.654, indicating increased accuracy from Block 1 (48.0%) to Block 2 (56.0%), and from Block 2 to Block 3 (58.5%), both *p* < 0.001 (with Bonferroni correction; Figure 2B, lower). The interaction between task and block did not reach significance, *F*(2, 30) = 2.40, *p* = 0.108.

The paired *t* test showed a drop in accuracy from the 6^th^ mini-block (56.8%) to 7^th^ mini-block (50.4%), *t*(15) = 2.11, *p* = 0.026 (one-sided), while the difference in accuracy between the 1^st^ mini-block (45.5%) and the 7^th^ mini-block was not reliable, *t*(15) = 1.86, *p* = 0.083 (Figure 3B, right). These results suggested that the improved performance in the naming task was not fully transferred to the discrimination task. In addition, there was no difference in accuracy between the two mini-blocks within Block 1, *t* < 1, suggesting the lack of transfer from trained viewpoints to untrained viewpoints.

For the persistence of the training effect, the ANOVA showed a significant main effect, *F*(2, 26) = 15.96, *p* < 0.001, *η*^2^ = 0.551, with the accuracies in the last training block (60.7%) and the post-test (56.3%) both higher than the accuracy in the first training block (47.1%), *p* < 0.001 and *p* = 0.015, respectively (with Bonferroni corrections) whereas the accuracy difference between the last training block and the post-test did not reach significance, *p* = 0.127. The BF analysis showed that B_0-_ = 0.288, which equals to B_-0_ = 3.469, indicating that the alternative hypothesis (‘the accuracy in the post-test was lower than the accuracy in the last training block’) was 3.469 times more likely to be true than the null hypothesis (‘the accuracy in the post-test was equal to or higher than the accuracy in the last training block’) (Figure 2B, lower). These results suggested a reduction of recognition performance after one week, but not to the initial pre-training level.

#### Comparisons across the four experiments

The above analysis showed consistent different patterns of training effect when the naming task was followed by the discrimination task (Experiments 1A and 2A) and when the task order was reversed (Experiments 1B and 2B), irrespective of whether the alternative choices increased over the 3 blocks (Experiment 1 vs. 2). To investigate whether the different patterns were due to different choice difficulties of the four experiments, we conducted a 2 (task type: naming vs. discrimination) * 2 (choice difficulty: 2 alternative choices vs. 4 alternative choices) ANOVA on the ACAs in the first mini-block. Only the ACAs in the first mini-block were included in the analysis in order to avoid any contamination from the improvement by training. As a result, neither the two main effects nor the interaction between the task type and choice difficulty reached significance, all *p* > 0.1, suggesting comparable starting capabilities in recognizing the simulated pictures regardless of the task type (naming vs. discrimination), and regardless of the alternative choices (2 vs. 4). As such, the observed different patterns cannot be reduced to be a by-product of the different tasks or different number of alternative choices.

To reveal which training procedure produced the best improvement in object recognition, we calculated the improvement by subtracting the ACAs in the first mini-block from the ACAs in the last mini-block, and conducted a 2 (task order: naming followed by discrimination vs. discrimination followed by naming) * 2 (choice difficulty: increasing alternative choices from 2 to 6 vs. constant 4-alternative choices) ANOVA on the ACAs difference. The results showed a trend of task order, *F*(1, 61) = 3.54, *p* = 0.065, *η*^2^ = 0.054, whereas the main effect of choice difficulty, *F*(1, 61) = 1.00, *p* = 0.321, and the interaction, *F* < 1, were not significant. These results suggested a better overall improvement with the training procedure where the naming task was followed by the discrimination task than the reverse order (18.2% vs. 14.8% in ACA).

### Discussion

In the present study, we simulated RI vision and investigated if object recognition in RI patients can be improved and maintained through training. Our results from two different tasks consistently showed improved recognition performance over two repetitions of the object pictures in both tasks, and the recognition performance persisted at least for a week. These results accord well with the literature on object perceptual learning (Bi and Fang, 2013; Fine and Jacobs, 2002) and provide important new evidence that the visual abilities of RI patients can be strengthened and maintained. Notably, the overall improvement (18.4% when the naming task was followed by the discrimination task) of familiar object recognition in RI vision through such a short period of training was already comparable to the improvement (around 20%) that healthy adults achieved over 5 days of training (Furmanski and Engel, 2000).

An important issue we focused in the present study was the generalization of the training effect. In particular, we investigated the transfer of object recognition to different task contexts, and to new viewpoints of the trained objects. As a result, we found that the improved recognition performance in the naming task persisted in the following discrimination task, suggesting true object recognition, independent of task demands. By contrast, the recognition performance improved by the discrimination task dropped in the following naming task, suggesting a lesser extent, if not the absence, of task transfer. In addition, the improved object recognition transferred to the untrained viewpoints of the trained objects in the naming task but not in the discrimination task. One plausible explanation might be that the naming task was easier than the discrimination task, thereby producing larger transfer (Ahissar and Hochstein, 1997). This account, however, can be ruled out because the initial performance in both tasks was comparable and the transfer patterns were consistent in both experiments regardless of task difficulty.

It has been suggested that task-independent plasticity reflects the changed representation of a feature or object in the visual cortex, in contrast to task-dependent plasticity that is associated with changes in the mechanism specific for the trained task, such as the attentional system (Watanabe and Sasaki, 2015). Previous perceptual learning studies with healthy adults do not always show task-independent plasticity, as there were failures in transferring the trained perceptual performance to a different task (Huang et al., 2007; Westheimer et al., 2001). For example, Huang and colleagues (2007) trained one group of healthy adults to detect a coherent motion signal (detection task) and the other group to discriminate the direction of the coherent motion signal (discrimination task). They found that the group trained with the detection task showed improvement in only detecting but not in discriminating the motion signal, whereas the group trained with the discrimination task showed improvement in both tasks. The asymmetry was explained as a result of task-relevancy in the way that detecting motion signal was necessary for discriminating the particular direction whereas the precise direction was unnecessary for detection and could be actively inhibited (Huang et al., 2007; Tsushima, Sasaki, and Watanabe, 2006). This finding suggested that the transfer of perceptual learning was dependent on the top-down attentional set in the training task. From this perspective, although there were improved performances in both tasks in the present study, the improvement in the naming task may be contributed mainly by the change in representations of the trained objects, whereas the improvement in the discrimination task may due to the processing strategy at hand.

It is well established that top-down information, such as task context, plays a critical role in object recognition (Bar, 2004; Trapp and Bar, 2015). Different task contexts engage different processing strategies, which in turn recruit distinct neural substrates to construct object representations in the brain (Op de Beeck and Becker 2010; Song et al., 2010; Wong et al., 2009a; 2009b). Two processing strategies in object recognition have been documented: a holistic processing strategy which treat the object as an undifferentiated representation, and a parts-based processing strategy which decompose the object into parts and the relations among parts (Farah, 1990). The holistic strategy was found to be dominant in an associative learning task of object recognition where subjects were trained to construct associations between the presented novel object and a particular word, while the part-based strategy was dominant in a discrimination learning task where subjects were trained to choose a target object out of other distracting objects (Song et al., 2010).

The naming task and the discrimination task in the present study was similar to the associative learning task and the discrimination task in Song et al. (2010), respectively. Accordingly, a holistic strategy was supposed to be engaged in the naming task while a parts-based strategy in the discrimination task, which in turn should have caused changes in the corresponding representation areas in the brain. However, unlike the unambiguous sensory inputs of the objects in the previous studies (Song et al., 2010; Wong et al., 2009b), the coarse sensory inputs of the object in the present study limited the parts-based processing and may encourage a holistic strategy in both tasks instead (Trapp and Bar, 2015). Therefore, only the naming task, but not the discrimination task, could benefit from the processing strategy and caused representation changes in the brain so as to produce the transfer effect.

In summary, we recommend the naming task as the training regime for RI patients to achieve persistent and flexible object recognition for the following three reasons: 1) the naming task produces higher overall improvement than the discrimination task in object recognition given the same amount of trainings; 2) the improvement in object recognition through the naming task can be generalized to discriminate different objects (i.e., the discrimination task); 3) the improvement in object recognition through the naming task can be generalized to new viewpoints of the trained objects; 4) the improved object recognition through the naming task fully persisted after one week. Our findings not only provide insights to the development of training protocols for RI patients, but also can help patients and their families to evaluate the expected effects before making the decision for or against implantation.

## Acknowledgments

This project was supported by the European Fund for Regional Development (EFRE, project number: ZS/2016/04/78113) from the Center for Behavioral Brain Sciences (CBBS). We thank Ariel Rokem and Michael Beyeler for their help in using the Pulse-to-percept software and Elisabeth Rosenblum for assisting in data collection.

---

**Supplementary Figure 1.**
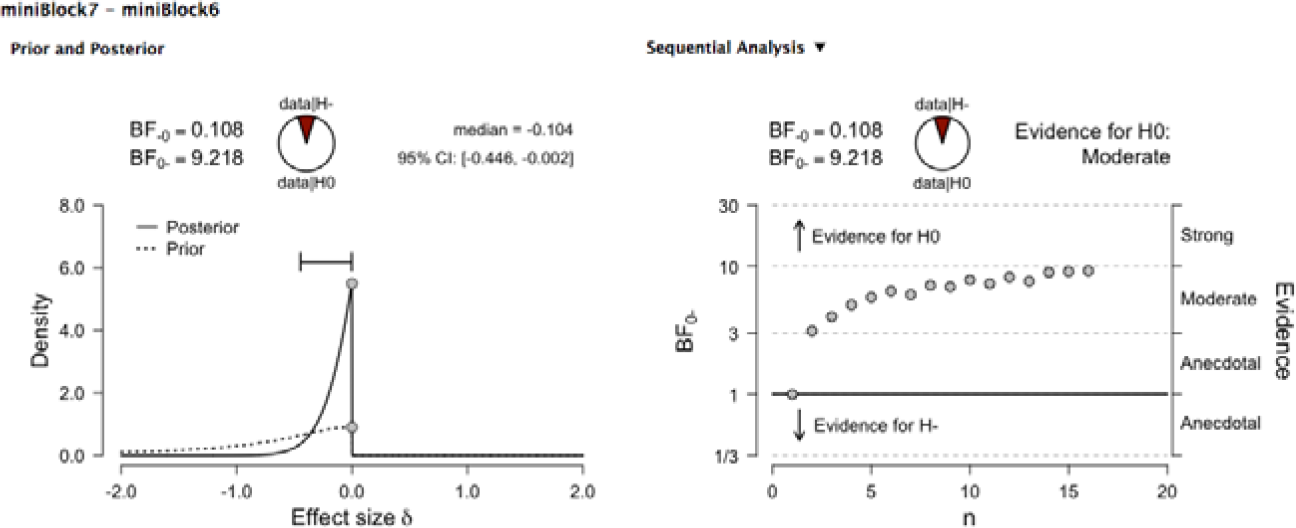
S1. An illustration of the one-sided Bayes factor analysis in JASP. Data from Experiment 2A (ACC_7th_ vs. ACC_6th_) are shown as an example. As a default setting in JASP, a Cauchy distribution with a scale of 0.7 was assumed as the prior (left panel). The ‘-’ in BF_-0_/BF_0-_ (i.e., the variant of BF_10_/BF_01_ for one-sided test) denotes the direction of the alternative hypothesis ‘ACC_7th_ < ACC_6th_’. The evidence for the null hypothesis increases as a function of the sample size (right panel)

